# Redefining and subtyping of major depression based on brain functional connectivity signatures with high generalization: an ensemble hybrid framework

**DOI:** 10.1101/2022.06.07.495073

**Authors:** Kaizhong Zheng, Hongbing Lu, Yuxia Wu, Huaning Wang, Dewen Hu, the DIRECT consortium, Badong Chen, Baojuan Li

## Abstract

Current clinical diagnosis of major depression is solely based on subjective symptoms and signs rather than the underlying biological mechanisms, which has hindered the improvement in treatment effectiveness and outcomes. Redefining and subtyping major depressive disorder (MDD) based on neural circuits has been an emerging consensus. Here, we proposed a novel ensemble hybrid deep-learning framework based on graph neural network for redefining and subtyping of MDD with the brain’s “fingerprinting”, i.e. nuanced functional connectivity profiles. A graph neural network (GNN) based supervised model was first employed to discriminate MDD patients from healthy controls. The graph embedding features obtained from the supervised classifier was then employed for subtyping of MDD with an unsupervised clustering by fast search and find of density peaks (CFDP) approach. The generalization accuracy and reproducibility of the proposed framework were further validated with one of the largest resting-state fMRI data set for MDD which included 2428 participants recruited from 25 independent sites. Our classifier showed a high generalization of 75% for leave-one-site-out cross validation. In addition, we obtained three biologically homogenous MDD subtypes characterized by impairments in subcortical-limbic, DMN-FPN, VN-SMN circuits, respectively. Notably, we demonstrated that subtyping of MDD according to graph embedding features showed high robustness and reproducibility across different sites. The proposed framework is not only essential for redefining and subtyping of MDD, but also paves the way towards precision medicine of mental disorders.

## Introduction

As a debilitating mental disorder, major depressive disorder (MDD) is regarded as the world’s most serious psychiatric disorder [1, 2] and represents the second leading contributor to chronic disease [3, 4]. However, the treatment efficacy is still unsatisfactory and responses have been divergent. Current diagnosis with the Diagnostic and Statistical Manual of Mental Disorders 5^th^ edition (DSM - 5) is based on subjective symptoms and signs rather than the underlying biological mechanisms. Clinically, an MDD diagnosis can be given when a patient shows at least five of nine symptoms [5], which have high heterogeneity within major depression and overlapping phenotypes across psychotic disorders [6–9], and has restricted progress towards improving treatment effectiveness and outcomes. Considering the above, a “precision medicine for psychiatry” project has launched[10]. Within the framework of the Research Domain Criteria (RDoC), mental disorders are suggested to be redefined as “disorders of brain circuits” and subtyped into biologically homogenous subgroups based on underlying genetic and neural mechanisms beyond observable symptoms [10].

Pioneering studies using biological and cognitive signatures are deconstructing traditional symptom-based categories to redefine and provide biologically meaningful subtypes of psychiatric disorders using the machine-learning method [11–15]. The functional connectome that is recognized as the fingerprint of the brain [16, 17] may serve as a potential candidate biomarker for mental disorders. Li et al. developed a framework for schizophrenia identification and subtyping based on functional striatal abnormalities [13]. As for subtyping of MDD, four neurophysiological subtypes that were predictable of treatment responses were identified according to functional connectivity profiles [5]. Although the promise of the machine-learning-based redefining framework has been widely appreciated, the generalization of the classifier which is considered as “a bare minimal requirement” [14] for its translation into clinical diagnosis and prognosis remained to be established. Validation of the generalizability of neuroimaging biomarkers to unseen data collected from completely different sites remains to be elusive [5, 18–20] due to the high biological heterogeneity of the disorders. Many frameworks have failed to demonstrate acceptable generalization for independent datasets collected at multiple sites [21] due to over-fitting and nuisance variables [14]. Until now, the highest generalizable accuracy of MDD classifiers reported in literatures was only 66% for an independent validation dataset from 5 different sites [20], hindering their possible application in clinics.

It has been suggested that ensemble hybrid frameworks combining supervised and unsupervised approaches could be employed to overcome the problems caused by heterogeneity and facilitate advance precision diagnostic [20, 21]. Thus, we proposed a novel ensemble hybrid deep-learning framework for brain network (EH-BrainNN) which adopted a “lumping” and “splitting” strategy for redefining and subtyping of MDD based on functional connectivity profiles. Firstly, we established a supervised approach to select a relatively small set of features that could robustly discriminate healthy controls and patients while lumping all subtypes of patients together. Graph neural network (GNN) which could capture the inherent topological structure of functional connectivity profiles was employed to identify a small set of explainable features. We then proposed a unsupervised clustering model to automatically identify biologically homogenous subtypes with the clustering by fast search and find of density peaks (CFDP) algorithm. Notably, the CFDP algorithm could identify the clusters regardless of their shape and the dimensionality of the space and select the correct number of clusters automatically [23].

The generalization accuracy and reproducibility of the proposed framework were further established with the REST-meta-MDD data set provided by the DIRECT consortium. This data set represents one of the largest MDD data set with fMRI images collected from 2428 participants including 1300 MDD patients and 1128 healthy controls (HCs) in 25 independent sites. For our supervised redefining model, the generalizability of the classifier was comprehensively validated by leave-one-site-out cross validation which achieved an average generalization accuracy of 75% over all the sites. In regarding to the unsupervised subtying model, the CFDP method robustly obtained 3 MDD subtypes. More importantly, the functional connectome signatures of each subtype also showed high consistency across different sites, indicating high reproducibility and generalization of the proposed ensemble framework. Thus, our study not only validated the feasibility of redefining and subtyping of mental disorders with underlying neural circuits measurements beyond symptoms, but also pave the way towards precision medicine of mental disorders.

## Results

### Overall framework of EH-BrainNN

The workflow of the proposed ensemble hybrid framework (EH-BrainNN) is shown in Fig.1. It is mainly consisted of three modules: a backbone prediction model, explanation generator and MDD subtypes generator (Fig. 1a). The backbone prediction model which included the Graph Isomorphism Network (GIN) [22] encoder and the bilinear mapping second order pooling was used to identify a robust signature from resting-state fMRI data that could discriminate depressed patients from healthy controls (Fig. 1b 2-3). The explanation generator funished interpretable explanations for classification results and explored the topological attributs of robust signature extracted from backbone prediction model (Fig. 1b 4). The MDD subtypes generator identified subtypes of MDD and investigated the association between patterns of subtypes and clinical symptoms (Fig. 1b 5).

**Fig. 1.**
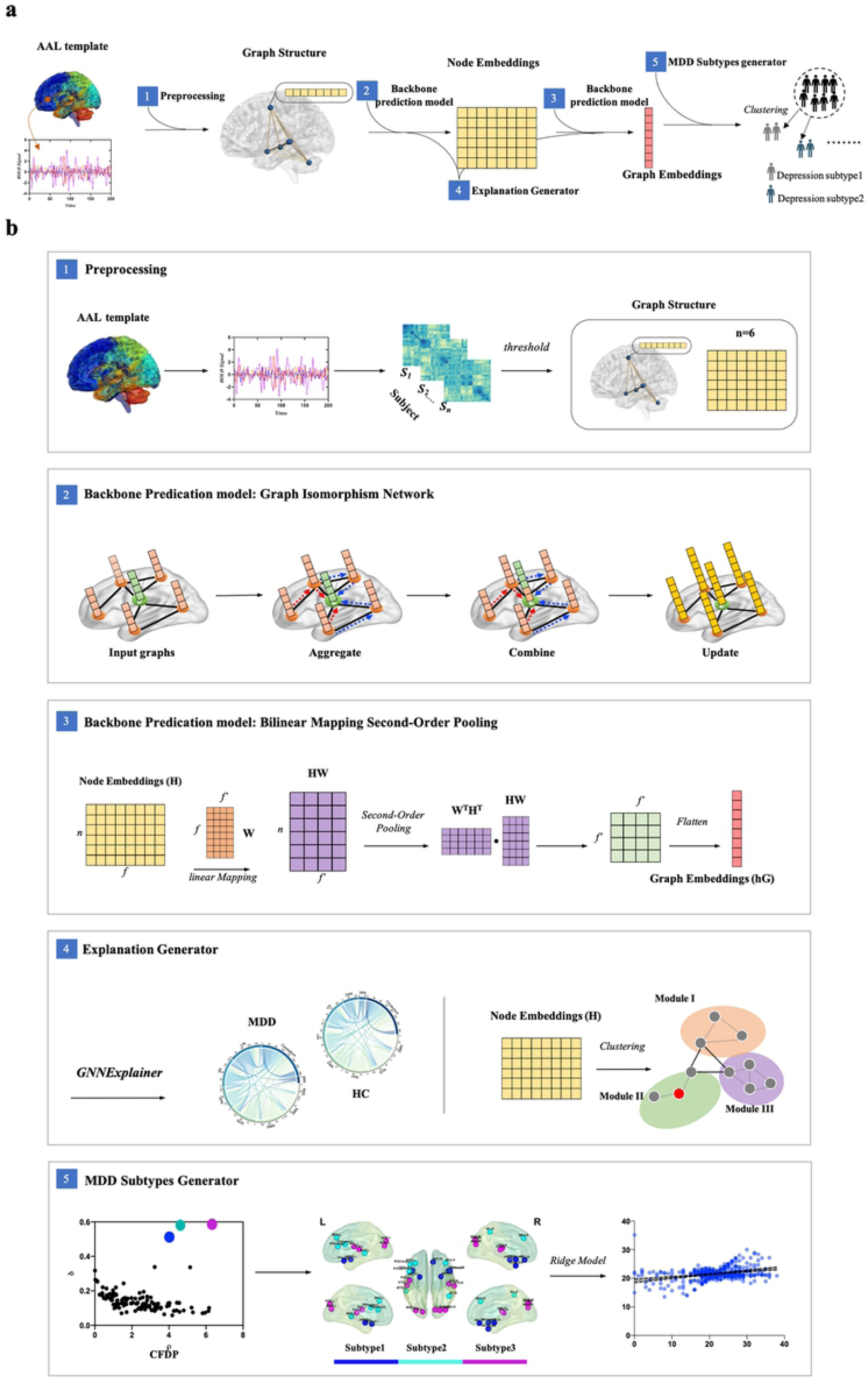
Overall ensemble hybrid frameworks combining supervised and unsupervised approaches (EH-BrainNN). **a** Overall ensemble hybrid framework based on graph deep-learning and data-driven clustering approaches for the identification of depression subtypes. **b** (1) Graph structure and the node feature were generated through preprocessing. (2) Schematic of the GIN model was mainly comprised of three steps: aggregating the information of neighbor nodes, constructing the node embeddings and updating the node embeddings. (3) Graph embeddings were constructed from node embeddings by the bilinear mapping second order pooling. (4) We used GNNExplainer model to make the interpretable explanations for classification results and used clustering using the clustering by fast search and find of density peaks (CFDP) to test the modularity of node embeddings. (5) CFDP was used to identify MDD subtypes. We further investigated the association between graph embeddings and clinical profiles for depression symptoms (HAMD) in each MDD subtype using leave-one-out ridge regression.

### Generalization Performance

#### Ten-fold cross-validation

To fit a model and optimize its hyperparameters, we conducted 10-fold cross-validation procedure [23]. Specifically, we divided the dataset into the training set (9 folds out of 10 folds), which was used for training a model, and the testing set (1 fold out of 10 folds) for testing a model. Using 10-fold cross-validation, EH-BrainNN achieved an accuracy of 69%. Figure 2a-b show sensitivity-specificity and precision-recall curves based on model predictions on the training data and testing data, respectively. The corresponding AUC value of the testing data in the sensitivity-specificity and precision-recall curves were 0.742 and 0.682, respectively, suggesting that the model had formidable discriminatory ability. Furthermore, we employed a framework of permutation tests to assess the generalization performance of the classifier. Fig. 2c shows the permutation distribution of the estimate using the EH-BrainNN. Results indicated that the classifier learned the relationship between the fMRI data and the labels with a probability of being wrong of <0.0001. In addition to the high classification performance of EH-BrainNN, the framework provided an information-condensed signature (graph embeddings) which yielded robust binary discrimination between depressed patients and healthy controls. Fig. 2d demonstrates node embeddings and graph embeddings for a subject, respectively. The distribution of graph embeddings was different in the patients with MDD compared with that of healthy controls as shown in Fig. 2e.

**Fig. 2.**
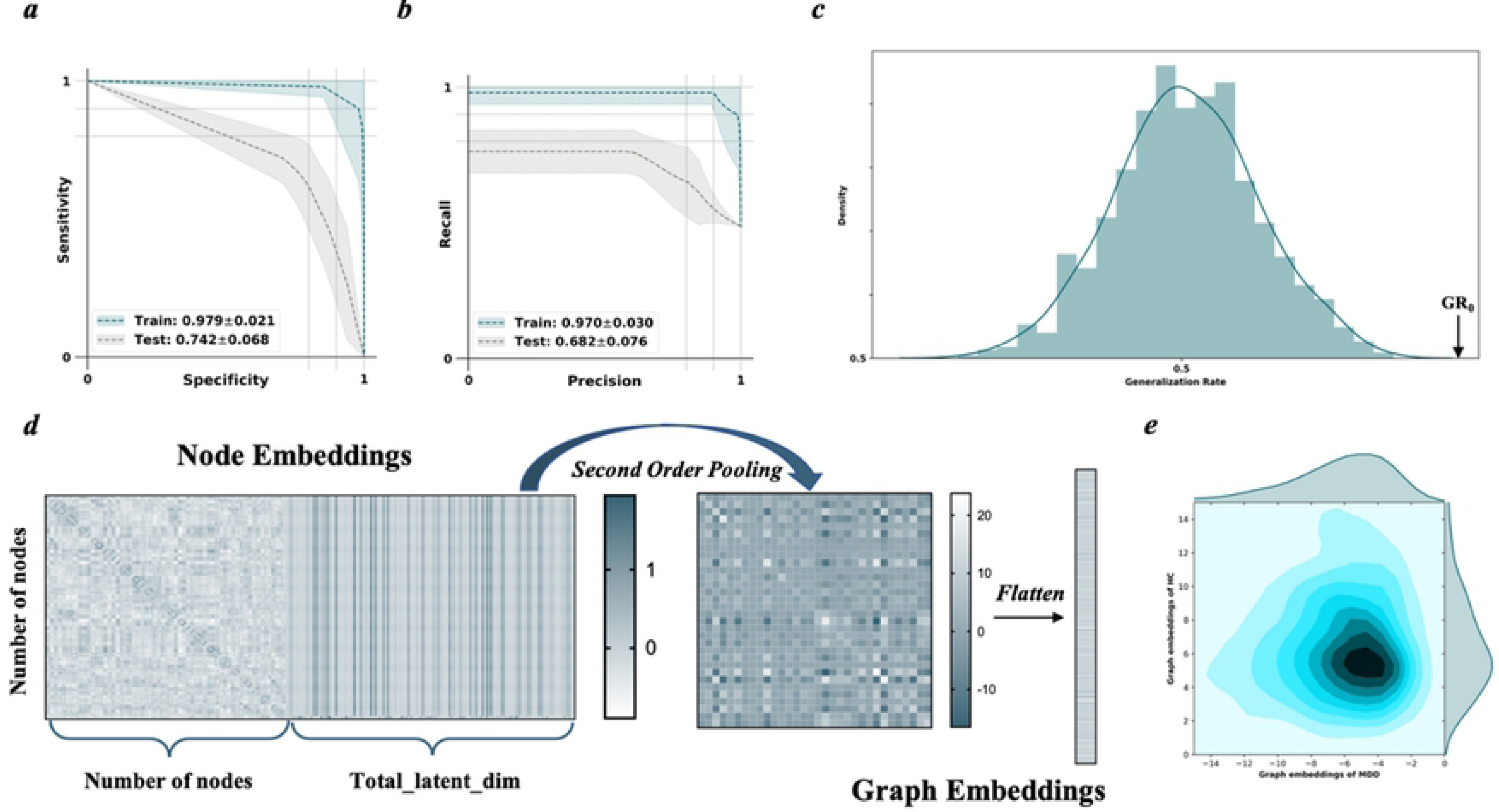
Performance of the GIN model for MDD’s disease classification using the 10-fold cross-validation procedure. **a-b** Sensitivity-specificity and precision-recall curves based on model predictions on the training data and testing data are depicted in dark green and gray, respectively. Meanwhile, the area under curve (AUC) values were computed for each sensitivity-specificity and precision-recall curve. **c** Permutation distribution of the estimate using the trained GIN model (repetition times: 1000) are shown, where *x*- and *y*-labels represent the generalization rate and probability density. GR_0_ denotes the generation rate gained by the GIN model trained on the real class labels. **d** Procedure of obtaining the graph embeddings from node embeddings in a single participant is depicted. Total_latent_dim = the number of nodes + the number of hidden units × (the number of GNN layers -1), where the number of hidden units and the number of GNN layers are the hyper-parameters and are set to 64 and 5. **e** The kernel distributions of graph embeddings of the MDD and HC populations are depicted.

Other metrics were used to further confirm the high classification performance of EH-BrainNN and robust discrimination of graph embeddings. We adopted *t*-distributed stochastic neighbor embedding (*t*-SNE) [24] using the function connectivity network and graph embeddings as inputs (Fig. 3) respectively. The *t*-SNE method is a visualization tool through transforming high-dimensional data into their low-dimensional representation of those data. The *t*-SNE plot of functional connectivity networks revealed no clear discrimination between depressed patients and healthy controls, while the *t*-SNE plot of graph embeddings resulted in diagnosis-specific clustering of the participants. These results suggested that the graph embeddings may serve as a robust neuroimaging biomarker in the patients with MDD.

**Fig. 3.**
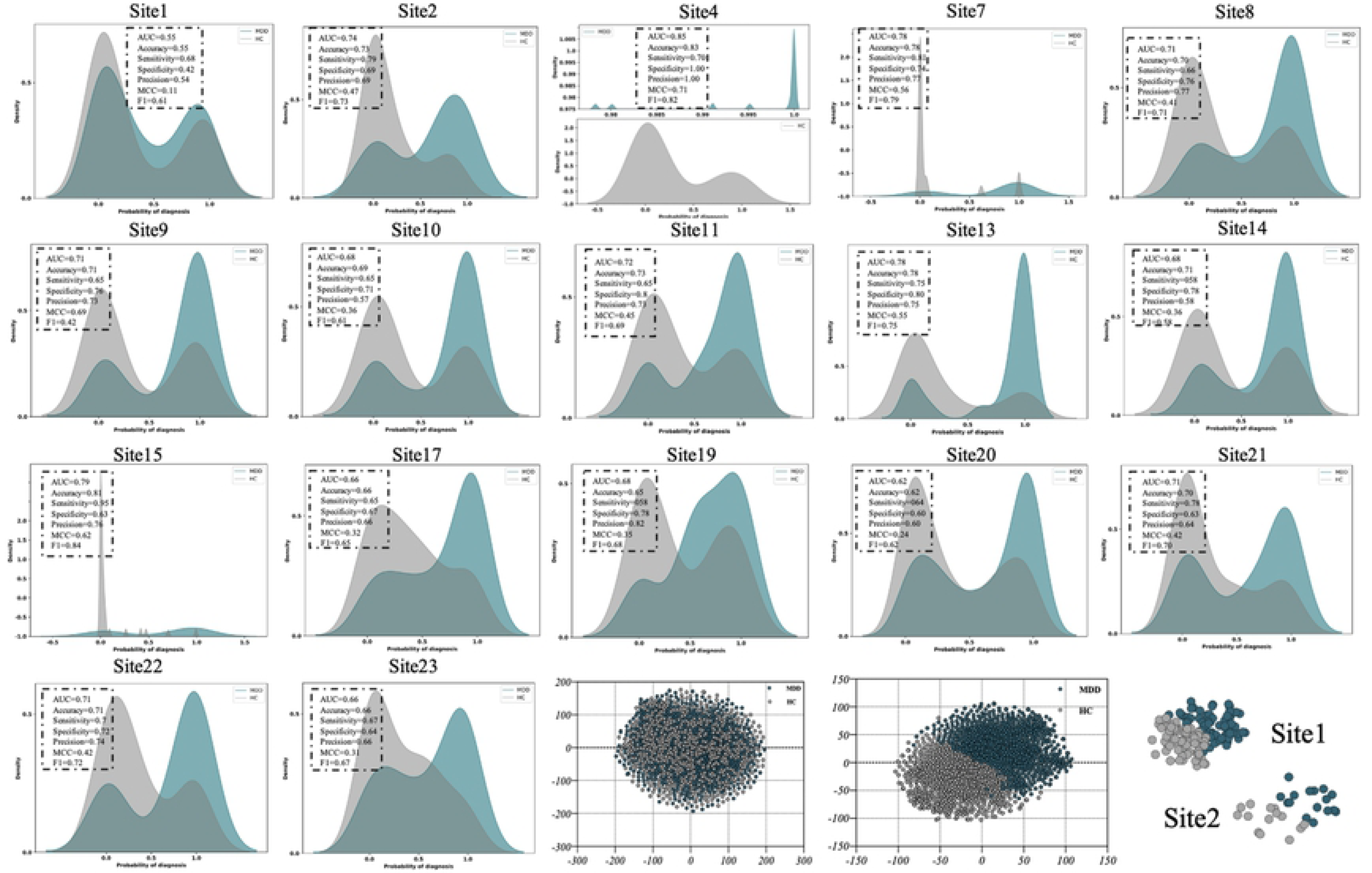
Generalization performances of the MDD classifier using a leave-one-site-out analysis. The probability for the diagnosis of MDD for each neuroimaging site between the patients with MDD and healthy controls are depicted in dark green and gray. In addition, the area under the curve (AUC), accuracy, sensitivity, specificity, precision, matthews correlation coefficient (MCC) and F1-score were also calculated. In addition, we show visualization of functional connectivity networks and graph embeddings using the t-SNE method. More details are described in the Supplementary Figure S5. AUC, the area under the curve; MCC, matthews correlation coefficient; MDD, major depressive disorder, HC, healthy control.

#### Leave-one site out cross-validation

To further assess the generalizability of classification model to unseen data collected at completely different sites, a leave-one-site-out analysis was performed by dividing the dataset into the training set (16 sites out of 17 sites) to train the model, and the testing set (the left site out of 17 sites) for testing a model (Fig. S3). A ComBat harmonization method [25–28] was applied to mitigate distinct site differences (*SI Appendix*, *Supplementary Methods*). Then we fitted a model to each site-sample while turning a model parameter in an inner loop of 10-fold cross validation, resulting 17 MDD classifiers. For each classifier, the classifier-output value (diagnostic probability) was considered to be indictive of an MDD diagnosis. Specifically, diagnostic probability of >0.5 was diagnosed as MDD, while diagnostic probability of <0.5 was diagnosed as HC [20]. We then calculated the area under the curve (AUC), accuracy, sensitivity, specificity, precision, Matthews correlation coefficient (MCC) and F1-score. Furthermore, we used *t*-SNE method to visual the graph embeddings from each site.

Fig. 3 shows the diagnostic probability density distributions of the MDD and HC populations for the 17 sites, indicating that 2 diagnostic probability density distributions were clearly separated by the 0.5 threshold for each site. To be specific, our classifiers achieved average generalization accuracy of 75% at 17 multiple completely different sites. Most classifiers achieved generalization accuracy of more than 70%, while the highest generalization accuracy was 83% (site 4). The AUC, sensitivity, specificity, precision, MCC and F1-value were 0.85, 70%, 100%, 100%, 0.71 and 0.82, respectively for the site 4. Furthermore, we found that the *t*-SNE plot of graph embeddings revealed clear discrimination between depressed patients and healthy controls for each site (Fig. S4). These observations indicated an advantage of our study that the EH-BrainNN was able to generalize well on the completely different multi-sites.

### Interpretability Analaysis

#### Modular analysis for node embeddings

To test whether node embeddings obtained from backbone prediction model are provided with known attributes of brain topology, we investigated the modularity of complex brain network through dividing the whole-brain network into distinct functional networks based on the node embeddings using clustering by fast search and find of density peaks (CFDP) [29]. The brain’s functional systems have the basic organizational characteristics of the complex networks such as small-world topology [30–33], highly connected hubs [34, 35] and modularity [36]. Modularity is supported by the social networks which represent some communities including networks of friendships or other acquaintances between individuals [36]. As same as the social networks, the brain’s functional systems consist of a number of functional modules. Each module encompasses several densely interconnected nodes, while there are relatively few connections between different modules. The results of modular analysis are shown in Fig. 4. All the 116 nodes were clustered into 3 modules in both groups (Fig. 4a-d), which clearly suggested that node embedding caputred the inherent topological information of functional brain networks. Fig. 4 e-f show that 12 nodes were clustered into different modules in the patients and healthy controls. The majority of these 12 nodes belong to the fronto-parietal network (FPN).

**Fig. 4.**
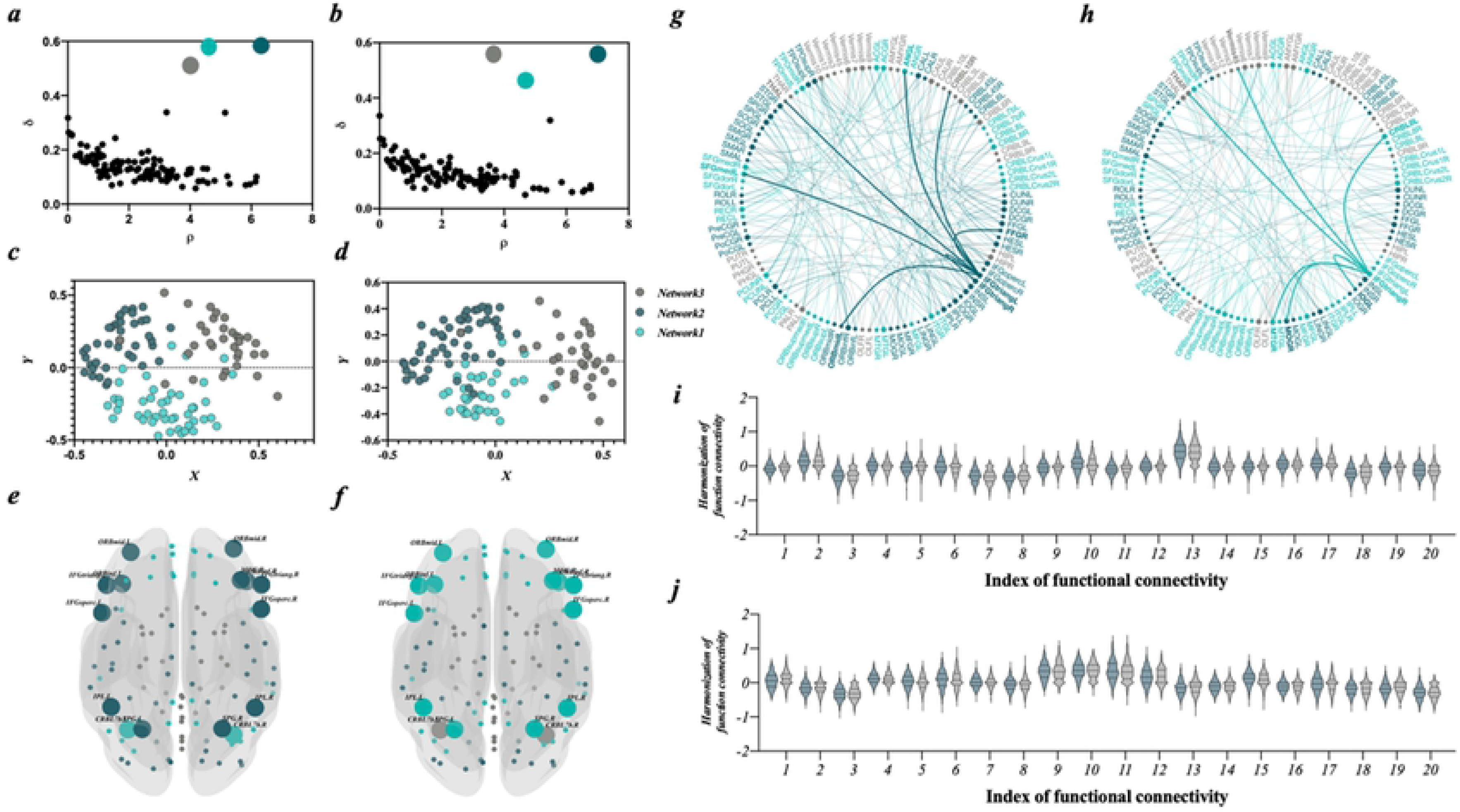
Interpretability Analaysis for MDD diagnosis. **a-f** the modularity of complex brain network was based on the node embeddings using the CFDP. **a-b** Corresponding decision graphs in both groups including patients with MDD and healthy controls respectively, with the centers colored by cluster. The light green represented the network1, the dark green represented the network2 and the gray represented the network3. **c-d** Node distributions for groups of MDD and HC. **e-f** 12 nodes assigned to different networks in both groups are depicted. **g-j** Interpretable explanations for classification results using GNNExplainer. **g-h** subgraphs of HCs and patients with MDD associated with MDD diagnosis are shown, respectively. **i-j** Reproducibility of important functional connections within subgraphs associated with MDD diagnosis. The dark green and gray represented MDD and HC respectively. Of node, these functional connections have been harmonized.

#### Interpretable explanations for classification results

To obtain important subgraph structure for an MDD diagnosis, we adopted GNNExplainer [37], model-agnostic approach for providing interpretable explanations for classification results. Briefly, we regarded GNNExplainer as an optimization task that maximized the correspondence between the classification results of backbone prediction model and distribution of possible subgraph structures. Unlike other traditional methods, GNNExplainer is able to obtain subgraphs of HCs and patients with MDD separately, which makes it easier to identify the similarities and differences between them (Fig 4). Fig. 4 g-h show subgraphs of HCs and patients with MDD associated with MDD diagnosis respectively and the most connected node and its edges are in bold. The most connected node belongs to fronto-parietal network, suggesting that fronto-parietal network played an essential role in diagnosis and network modularity. Fig. S6 A-B show subgraphs in patients with MDD and healthy controls. We observed that common subgraph in patients and healthy controls was composed of functional connectivity between subcortical network and limbic network, between limbic network and visual network, and between cerebellum and other functional networks. By contrast, distinct subgraph between the two subject groups was composed of functional connectivity between default mode network and subcortical network, limbic network, somatomotor network; and functional connectivity between fronto-parietal network and visual network, limbic network. In particular, limbic network and fronto-parietal network exhibited the high discriminative power between the MDD and HC subgraph. A detailed list of the subgraphs is provided in Tab. S5-8.

### Subtyping Analysis

#### Graph embeddings define three depression subtypes

According to our overall frameworks, we further adopted unsupervised learning to these high-dimensional signatures obtained from deep-learning model to identify subtypes of MDD. To ensure that cluster discovery was not confounded by the data distribution and the number of clusters, we proposed a data-driven method which automatically idenfied the number of clusters and MDD subtypes with graph embedding obtained from the backbone prediction model. This approach identified three depression subtypes which comprised 64.0%, 16.2% and 19.8% of the 828 patients with major depression, respectively (Fig. S7).

To illustrate specific decificts in each subtype of MDD, we compared the node embeddings between patients of each subtype with that of the healthy controls. The results showed distinct patterns of impairments in functional connectivity in different subtypes of MDD (Fig. 5). Specifically, MDD subtype1 was characterized by abnormal node embedding in subcortical network and limbic network. MDD patients of subtype 2 demonstrated impairments in a network mainly encompassed the default mode network (DMN) and FPN. Patients of subtype 3 mainly showed abnormal node embedding in the visual network (VN) and the Sensorimotor network (SMN).

**Fig. 5.**
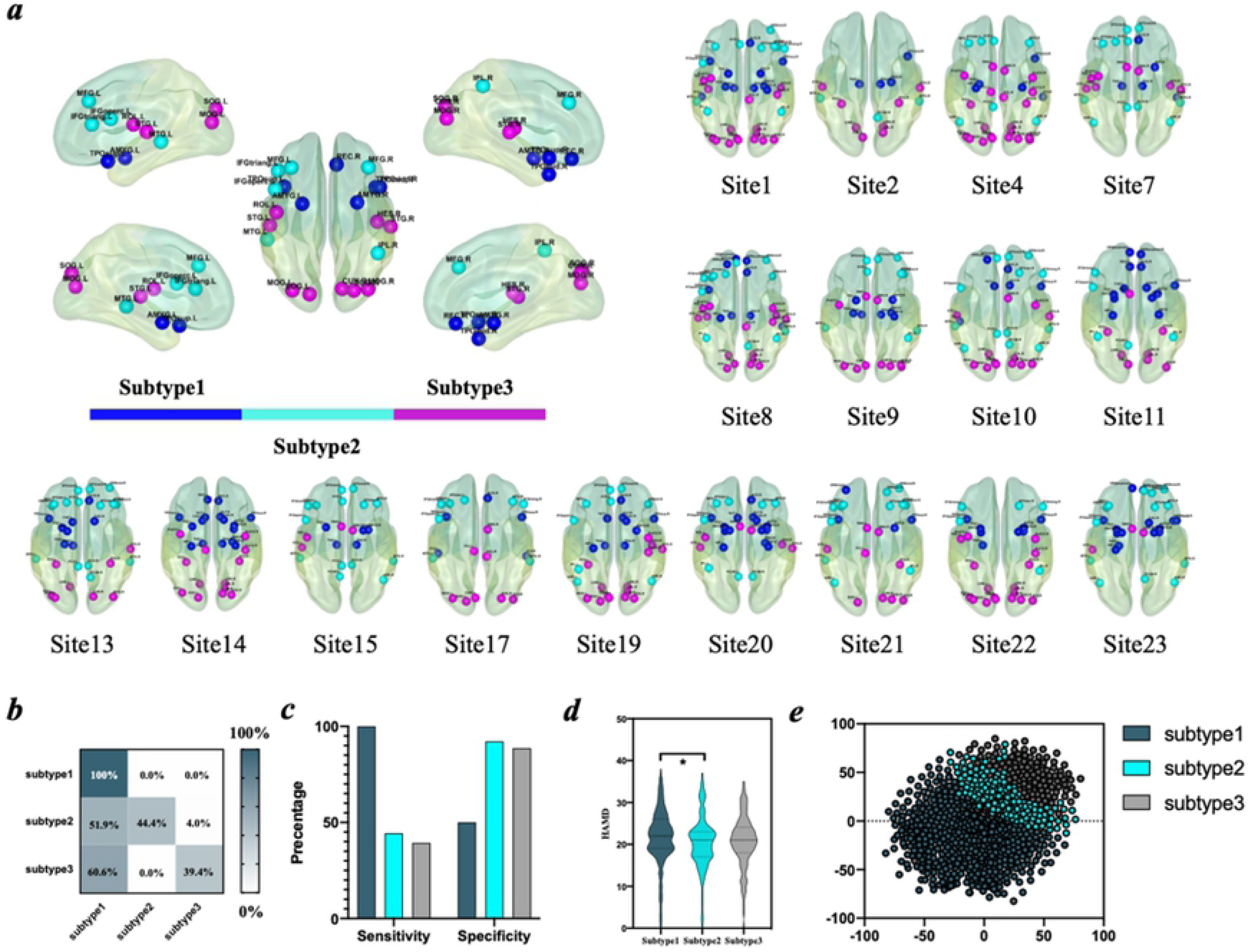
Graph embeddings define depression subtypes. **a** Neuroanatomical distribution of the abnormal patterns of node embeddings in three depression subtypes compared with HC using Wilcoxon rank-sum tests in all patients and 17 different sites. Different colors of ROIs define distinct depression subtypes. **b** Confusion matrix of accuracy by three depression subtypes for the classifier. **c** Sensitivity and specificity by three depression subtypes for the classifier. **d** A comparison of HAMD among three depression subtypes. **e** Graph embeddings from depression subtypes classifiers were used as inputs and a two-dimensional plot was generated using the *t*-SNE, where the dark green represented the depression subtype1, the light blue represented the depression subtype2 and the gray represented the depression subtype3.

We further validated the consistency and reproducibility of our identified MDD subtypes in each of the 17 independent site. Interestingly, we consistently identified three MDD subtypes in different sites. More importantly, functional connectivity patterns of the three subtypes were also highly reproducible across all the 17 sites (Fig. 5a).

In addition, we further developed classifiers for depression subtypes and adopted 10-fold cross validation to test the classification performance. Our classifiers achieved overall accuracy rates of up to 66.7% for three depression subtypes, and individual patient with different depression subtypes were identified correctly with sensitivities of 100%, 44.4%, 39.4% and specificities of 50%, 92.2%, 88.7% (Fig. 5 b-c). Furthermore, we adopted the *t*-SNE approach using the graph embeddings (Fig. 5 e) to test the performance of classifier. We found that the *t*-SNE plot of graph embeddings revealed clear discrimination among patients with three depression subtypes.

#### Correspondences between functional signature and MDD symptoms

To further investigate the association between patterns of discriminative functional signature and clinical symptoms, we used graph embeddings and clinical profiles for depression symptoms (HAMD) to generate GE-HAMD models using leave-one-out ridge regression. Fig. 6 shows the accuracy and error of GE-HAMD model predictions in each domain. The scatter plots show predicted and true clinical measures used to determine prediction accuracy (*r^2^*) for each participant (Fig. 6 b-c). We found significantly different prediction effect for the distinct clinical profiles in three depression subtypes. For example, the symptoms profiles of agitation were better predicted by graph embeddings of depression subtype 3. Next, prediction accuracy of the GE-HAMD models was further compared by a two-tailed Wilcoxon rank-sum tests of prediction errors (Fig. 6d). Five domains demonstrated significant differences among GE-HAMD models in three depression subtypes. Insight symptoms were better predicted by graph embeddings of subtype1 than subtype2 and subtype3 (subtype1 vs. subtype2: *p* = 0.0084; subtype1 vs. subtype3: *p* = 0.0312); Retardation scores were better predicted by graph embeddings of subtype2 than subtype1 and subtype3 (subtype1 vs. subtype2: *p* = 0.0276; subtype2 vs. subtype3: *p* = 0.0106); whereas, physical anxiety subtype1 vs. subtype3: *p* = 0.0290), hypochondria (subtype1 vs. subtype3: *p* = 0.0060) and weight loss (subtype1 vs. subtype3: *p* = 0.0019; subtype2 vs. subtype3: *p* <0.0001) were better predicted by graph embeddings of subtype3 than subtype1 and subtype2.

**Fig. 6.**
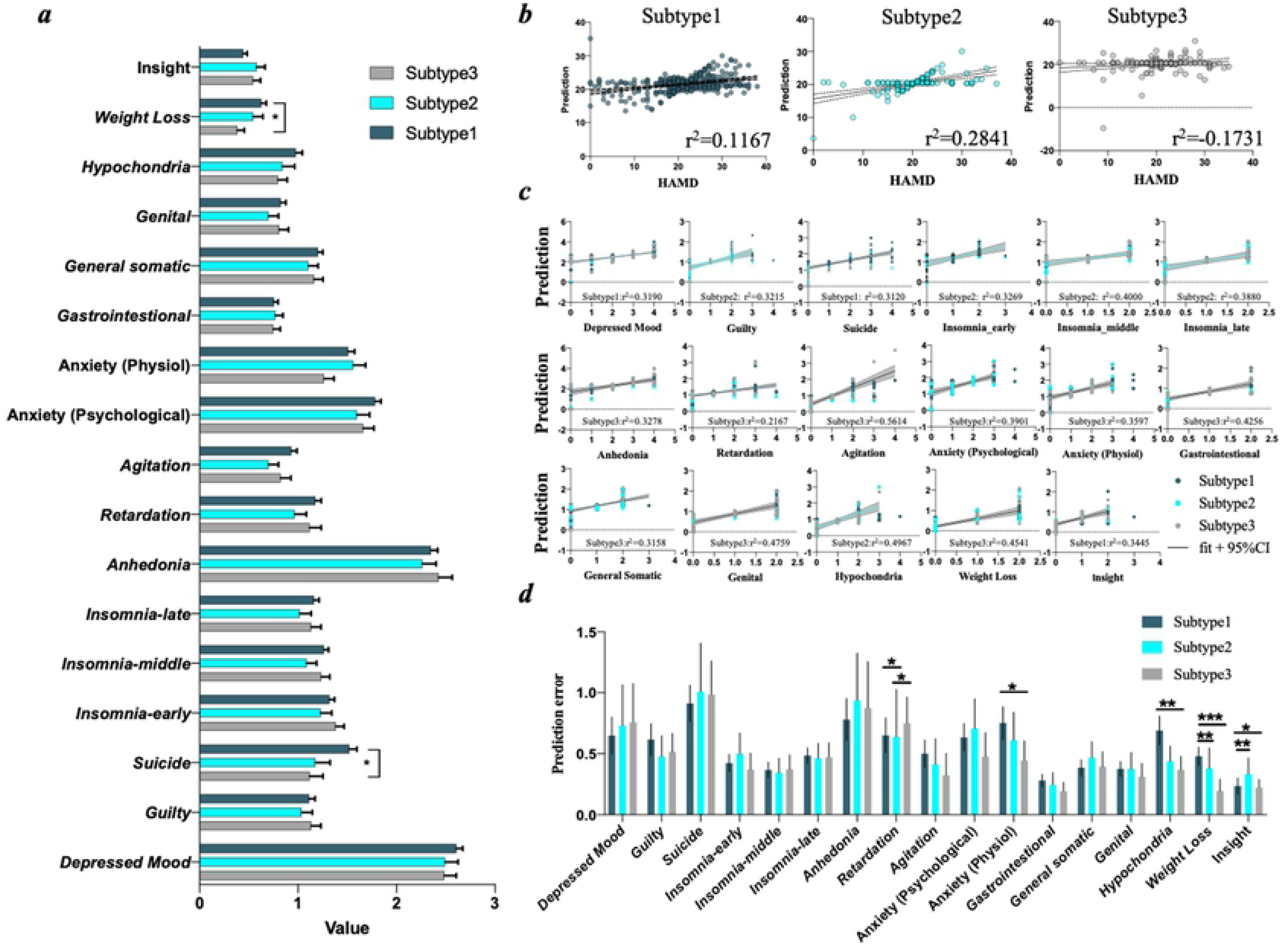
Subtype-specific clinical profiles for depression symptoms. **a** Comparison of clinical profiles for depression symptoms among three depression subtypes using Wilcoxon rank-sum tests. The scatter plots show the comparison between predicted and measured scores from GE-HAMD models for **b** HAMD and **c** clinical profiles for depression symptoms. **d** The bar graph shows prediction accuracy (the coefficient of determination, *r^2^*) across the 17 depressive domains in three depression subtypes.

### Effects of harmonization

To assess effects of Combat harmonization method, we repeated leave-one-site-out analysis without harmonization and compared prediction performances (Tab. S8). As a result, we found significant improvements in the prediction performances using harmonization for most sites. For example, the classifier for site 13 with harmonization achieved generalization accuracy of 78%, while the classifier for site 13 without harmonization achieved generalization accuracy of 64% merely. However, there were still no significant improvements in the prediction performances by harmonization for some sites such as site 4, which might be associated with the size and data distributions of the site and disease difference etc. Interestingly, we observed significant improvements in the prediction performances after harmonization for the sites with accuracy of less than 70%, but not after harmonization for the sites with accuracy of more than 70%. This suggested that the EH-BrainNN was able to obtain more robust and stable classification performance from different sites by the Combat harmonization.

## Discussion

In the current study, we established EH-BrainNN, a novel ensemble hybrid framework based on graph deep-learning and a data-driven clustering approach. With this framework, we first leveraged supervised learning method to develop a classifier for MDD which achieved a high average generalization accuracy of 75% at 17 multiple completely different sites for leave-one-site-out cross validation. These results suggested that graph embedding features constituted a “fingerprinting” of MDD which could reliably discriminate patients from HCs. After redefining MDD with this “lumping” supervised method, we then employed unsupervised CFDP clustering algorithm to obtain three biological homogenous subtypes of MDD. Importantly, the subtyping based on resting-state functional signatures demonstrated high robustness and reproducibility across all 17 different sites. Our results showed that the ensemble hybrid framework we developed for redefining and subtyping of MDD had high reproducibility and generalization which was crucial to ensure its translation into clinical diagnosis and prognosis.

The current study could largely promote the recent attempts of MDD redifing and subtyping according to neural mechanisms beyond observable symptoms by providng a novel ensemble hybrid deep-learning framework. Deep-learning approach serves as an extension of traditional machine-learning approach[38] and can automatically learn a latent high-level information from raw input data, making this method ideally suited to studying complex, subtle and scattered brain patterns [39, 40]. However, the potential advantages of deep-learning methods have not been fully exploited in neuroimaging domains. Functional connectivity profiles represent naunced structures of functionally-connected brain networks that were overlooked by previous studies that simply flattened functional connectivity profiles into a vector. To tackle this problem, we adopted GNN which represented functional connectivity profiles as graphs, thus could reserve the inherent structure of functional networks. In addition, although some previous attempts have been made to employ deep learning studies in discriminating patients with mental disorders and healthy controls [41–44], these models are based on ‘black-box’ algorithms which prevent the identification of underlying diagnostic decisions [45, 46]. We utilized the GNNExplainer which furnished indenfication of the structure of subgraphs that were crucial for diagnostic decisions. In order to obtain reliable and reproducible subtypes of MDD, we developed a unsupervised data-driven method which could automatically identifiy number and patterns of MDD subtypes by employing the CFDP algorithm.

Our classifiers achieved an average generalization accuracy of 75% for leave-one-site-out cross validation with a multicenter dataset collected from17 independent sites. Recently, there has been increasing attention on improving the generalization of the classifier. It is an emerging consensus that the generalization of a machine learning framework should be first proved before it can be considered as practical in clinical applications to ensure its reproducibility [14, 20]. The validation of the generalization for a classifier was even considered as “a bare minimal requirement” for its clinical application [14]. More importantly, the generalization should be tested with independent datasets from multiple sites [20]. Frameworks that showed high generalization on the diagnosis of ASD and Alzheimer’s disease have been established [45, 47, 48]. As for MDD, pioneering work by Yamashita and colleagues has attempted to develop an MDD classifier using logistic regression. They achieved an average accuracy of 66% with an external dataset which included data from 5 different sites [20]. In the current study, we used a leave-one-site-out cross validation to comprehensively test the generalization of our algorithm with one of the largest MDD dataset which recruited 1604 patients with MDD from 17 sites. We obtained an average generalization accuracy of 75%, suggesting that the classifier we established based on FC profiles and GNN has high scientific reproducibility and acceptable clinical applicability. As an objective neuroimaging biomarker, the graph embedding features could thus help with the redefining of MDD. The classifier we established could provide clear diagnosis boundary which allows accurate diagnosis and effective intervention of MDD.

Another major contribution of the current study is that we consistently identified three depression subtypes across all the sites. It is well known that MDD is not a unitary disease but a heterogeneous disorder since the patients present varied symptoms and respond divergently to treatment [5], suggesting that different biological mechanisms underline different subtypes. The importance of identifying diagnostic biomarker for MDD subtypes have long been recognized, by the solution is still unclear. Few attempts have been made while mixed results were reported. Drysdale and co-authors proposed a framework to identify MDD subtypes according to resting-state functional connectivity profiles based on canonical correlation analysis (CCA) and hierarchical clustering approaches. They found four neurophysiological subtypes using canonical correlation analysis (CCA) and hierarchical clustering approaches [5]. However, another recent study only identified 2 depression subtypes with CCA and K-means clustering algorithm [49]. It remains unaddressed whether consistent and reproducible MDD subtypes could be derived from different sites in order to provide clinically applicable biomarkers. To test this hypothesis, we adopt the data-driven CFPN algorithm which automatically obtain the number of MDD subtypes. Intriguingly, we consistently found three MDD subtypes across all the 17 sites. In addition, the functional connectivity signatures of the three subtypes were also shown to be highly reproducible. Patients of subtype1 mainly demonstrated impairments in a brain system encompassing subcortical network and limbic network. Patients of subtype 2 on the other hand were associated with deficits in the DMN and FPN. The third subtype were mainly associated with the SMN and VN. Our findings were in line with previous study by Drysdale[5], where subtype1, 3 and 4 showed abnormal connectivity within FPN and subcortical network and subtype1 and 2 were associated with impairments in limbic network. In addition, similar deficits in DMN, FPN, subcortical network and visual network were observed in subtype2 from previous study by Wang et al. [49]. These results suggested that patients with MDD as a group share some traits and common features of subtypes, which have potential to obtain biomarkers used for clinical guidance. However, these subtypes demonstrated overlapping without clear distinction unlike our depression subtypes in two previous studies. The current study thus extended previous studies and showed that highly consistent and reproducible MDD subtypes could be obtained with a large multisite dataset. Our findings could promote the development of personalized diagnostic and treatment strategy.

One limitation of the current study is that we were not able to investigate whether patients of different subtypes would response differently to antidepressant treatment since the Meta-MDD dataset only provided baseline clinical and neuroimaging data. We will further address this issue by applying rTMS to different subtypes and access the treatment efficacy. In addition, it will also be very important to develop precise targeting strategy and apply rTMS to subtype-specific target areas to ensure optimized treatment outcomes. Moreover, our initial analyses have showed that overlapping but unconformable correspondence between major depression and autism. We may further investigate whether there exists a “continuum” for mental disorders by including data from other disorders like schizophrenia, bipolar disorder, anxiety disorder, etc.in future work.

## Materials and methods

### Participants

The fMRI data was used from the DIRECT consortium which was lauched in 2017 aiming to pooling neuroimaging data collected from multiple indenpent sites to boost the statistical power of data analysis and finally facilitate the clinical application of findings from neuroimaging studies. DIRECT consortium provided one of the largest MDD dataset (http://rfmri.org/REST-meta-MDD). Invesigators from more than twenty-five research groups including our own had shared resting-state fMRI data. In the current study, 1604 participants (848 MDDs and 794 NCs) were selected according to exclusion criteria from a previous study [50] (*SI Appendix*, Fig. S1). Exclusion criteria mainly included incomplete information, bad spatial normalization, bad coverage, excessive head motion, and sites with fewer than 10 subjects in either group. In addition, three patients and two controls were removed from the sample due to incomplete time series data. Demographic and clinical characteristics of patients with MDD and HC subjects are provided in *SI Appendi*x, Tab. S1 and Fig. S2.

### Preprocessing

Our further analyses were based on preprocessed data. Preprocessing included discarding the initial 10 volumes, slice-timing correction, head motion correction, space normalization and temporal bandpass filtering (0.01-0.1 HZ). Statistical corrections removed effects of head motion, global signal, white matter and cerebrospinal fluid signals, as well as linear trends. After fMRI preprocessing, the brain was parcellated into 116 ROIs (regions of interest) according to the AAL (automated anatomical labelling) atlas [51]. The mean time courses of all the 116 ROIs were then extracted. We evaluated functional connectivity (Fisher’s r-to-z transformed Pearson’s correlation) between all ROI pairs. Therefore, a 116×116 symmetric matrix (functional connectivity network) *F* was obtained for each subject. Subsequently, we generated the graph structure and the node feature from functional connectivity network which was then provided as the input of graph isomorphism network.

### EH-BrainNN

We proposed a novel framework, i.e. EH-BrainNN for redefining and subtying of MDD. The EH-BrainNN included three core models as shown in Fig.1. The backbone prediction model and explaination generator were developed to redefine MDD with high generalization and the MDD subtypes generator was established to automatically identify MDD subtypes according to graph embedding signatures.

### Backbone prediction model

#### Model Procedure

Backbone prediction model was mainly consisted of Graph Isomorphism Network (GIN) [22] encoder and the bilinear mapping second order pooling. GIN which was one of the Graph Neural Networks (GNNs), was able to distinguish distinct graph structures effectively and achieves state-of-the-art performance on many graph classification benchmarks [22]. The GIN model mainly included three steps: first, given inputs, the GIN model computes neural messages between every pair of nodes. Second, for each node, the GIN model aggregates the representations of its neighbors. Third, the GIN model obtains the aggregated messages and constructs the new node embeddings. After obtaining the node embedding, we used the bilinear mapping second order pooling to generate the graph embedding. The graph embedding was employed to classify patients with depression from healthy controls. Mathematics description and performance measures are provided in *SI Appendix*, *Supplementary Methods*.

#### GIN model configurations

The GIN model consisted of 5 GNN layers (including the input layer) and each multi-layer perception (MLP) had two layers comprising a single hidden layer and an output layer. Each hidden layer was followed by batch normalization. All MLP also included non-linear operators such as ReLu. We used the Adam optimizer with a learning rate of 0.001 and the mini-batch size of 64. Dropout was used in the final dense layer.

#### Training, testing and hyperparameter optimization

Following the same training process [22], we performed 10-fold cross-validation strategy [23] to split out the training and testing sets from dataset and reported the average and standard deviation of validation AUC across the 10 folds within the cross-validation. In this procedure, the dataset was divided the training set (9 folds out of 10 folds) for training a model and the testing set (1 fold out of 10 folds) for testing the model. In addition, the hyper-parameters we tuned for the training dataset were: (1) the number of hidden units; (2) the batch size; (3) the number of total epochs; (4) the dropout ratio; (5) the number of iterations per each epoch. After debugging, all the hyper-parameters were fixed: hidden units of size 128, batch of size 32, the number of epochs 350, dropout ratio 0.5 and the number of iterations number 50.

### Explaination Generator

#### GNNExplainer to obtain brain markers elucidating the underlying diagnostic decisions

When the MDD classifier was generated, GNNExplainer [37] was employed to provide interpretable explanations for classification results. GNNExplainer is able to capture a compact subgraph structure and a small subset of node features that play a crucial role in GNN’s prediction. Mathematics description are provided in *SI Appendix*, *Supplementary Methods*.

#### Modular analysis based on node embeddings

Recent study has been verified that the node embeddings were provide with important topological information [52]. Therefore, we further investigated the modular property from the node embeddings using clustering by fast search and find of density peaks (CFDP) [29] (*SI Appendix*, *Supplementary Methods*).

### MDD subtypes Generator

#### CFDP to identify the subtypes of MDD

We proposed a unsupervised model with the clustering by fast search and find of density peaks (CFDP) [29] to automatically identify the depression subtypes using the graph embedding signatures. This clustering method is able to identify the clusters regardless of their shape and of the dimensionality of the space and automatically detect the optimal number of clusters unlike traditional clustering methods such as K-medoids method [53] which requires manually selection of cluster number. For the clustering procedure, we first computed the Pearson correlation *J_i,j_*between graph embeddings of every two participants. Then a distance *d_i,j_* between every pair of participants was defined as 1-*J_i,j_*. Finally, we used CFDP method using the *d_i,j_* as the input to obtain the depression subtypes. After identifying the depression subtypes, we used Wilcoxon rank-sum tests to compare patterns of node embeddings shared by all three depression subtypes and healthy controls. In order to further validate that the final three depression subtypes were indeed reflective of internally neurobiologically meaningful phenotypes related to brain patterns, we developed a GIN-based classifier to distinguish three depression subtypes using graph embeddings features. A 10-fold cross-validation was performed to measure the classification performance including accuracy, sensitivity and specificity.

#### Ridge Regression to investigate the brain-symptoms associations in different subtypes

Brain-symptoms associations were investigated using leave-one-out ridge regression models (Fig. 5). On the brain side, graph embeddings for each patient were used to predict clinical symptoms. Principal component analysis (PCA) was used to reduce the dimension of graph embeddings and components which explained 95% of variance were retained. On the symptoms side, there were 18 distinct clinical profiles for depression symptoms. For each ridge regression analysis, we built only one model to examine the relationship between graph embeddings and one clinical symptom score. In each loop, we optimized the regularization coefficient lambda to minimize prediction error through identifying the value of lambda between 1 and 10^4^. Performance measures of each model was provided in *SI Appendix*, *Supplementary Methods*. In addition, difference in prediction accuracy between graph embeddings and clinical symptom scores were measured by a two-tailed Wilcoxon rank–sum test on the prediction error (*y_prediction_*-*y_true_*)^2^.

## Acknowledgements

The authors thank the DIRECT consortium for providing access to the data.

## Funding information

This work was supported by the National Natural Science Foundation of China (62088102, U21A20485, 81974215, 81371478).

## Competing interests

All authors declare that they have no competing interests.

## Notes

### Competing Interest Statement

The authors have declared no competing interest.

## References

1. Ferrari AJ, Charlson FJ, Norman RE, Patten SB, Freedman G, Murray CJ, et al. Burden of depressive disorders by country, sex, age, and year: findings from the global burden of disease study 2010. PLoS Med. 2013;10(11):e1001547. Epub 2013/11/14. doi: 10.1371/journal.pmed.1001547. PubMed PMID: 24223526; PubMed Central PMCID: PMCPMC3818162.

2. DALYs GBD, Collaborators H. Global, regional, and national disability-adjusted life-years (DALYs) for 333 diseases and injuries and healthy life expectancy (HALE) for 195 countries and territories, 1990-2016: a systematic analysis for the Global Burden of Disease Study 2016. Lancet. 2017;390(10100):1260–344. Epub 2017/09/19. doi: 10.1016/S0140-6736(17)32130-X. PubMed PMID: 28919118; PubMed Central PMCID: PMCPMC5605707.

3. Otte C, Gold SM, Penninx BW, Pariante CM, Etkin A, Fava M, et al. Major depressive disorder. Nat Rev Dis Primers. 2016;2:16065. Epub 2016/09/16. doi: 10.1038/nrdp.2016.65. PubMed PMID: 27629598.

4. Belmaker RH, Agam G. Major depressive disorder. N Engl J Med. 2008;358(1):55–68. Epub 2008/01/04. doi: 10.1056/NEJMra073096. PubMed PMID: 18172175.

5. Drysdale AT, Grosenick L, Downar J, Dunlop K, Mansouri F, Meng Y, et al. Resting-state connectivity biomarkers define neurophysiological subtypes of depression. Nat Med. 2017;23(1):28–38. Epub 2016/12/06. doi: 10.1038/nm.4246. PubMed PMID: 27918562; PubMed Central PMCID: PMCPMC5624035.

6. Goodkind M, Eickhoff SB, Oathes DJ, Jiang Y, Chang A, Jones-Hagata LB, et al. Identification of a common neurobiological substrate for mental illness. JAMA Psychiatry. 2015;72(4):305–15. Epub 2015/02/05. doi: 10.1001/jamapsychiatry.2014.2206. PubMed PMID: 25651064; PubMed Central PMCID: PMCPMC4791058.

7. Jacobi F, Wittchen HU, Holting C, Höfler M, Pfister H, Müller N, et al. Prevalence, co-morbidity and correlates of mental disorders in the general population: results from the German Health Interview and Examination Survey (GHS). Psychol Med. 2004;34(4):597–611. Epub 2004/04/22. doi: 10.1017/s0033291703001399. PubMed PMID: 15099415.

8. Lee SH, Ripke S, Neale BM, Faraone SV, Purcell SM, Perlis RH, et al. Genetic relationship between five psychiatric disorders estimated from genome-wide SNPs. Nat Genet. 2013;45(9):984–94. Epub 2013/08/13. doi: 10.1038/ng.2711. PubMed PMID: 23933821; PubMed Central PMCID: PMCPMC3800159.

9. McTeague LM, Huemer J, Carreon DM, Jiang Y, Eickhoff SB, Etkin A. Identification of Common Neural Circuit Disruptions in Cognitive Control Across Psychiatric Disorders. Am J Psychiatry. 2017;174(7):676–85. Epub 2017/03/23. doi: 10.1176/appi.ajp.2017.16040400. PubMed PMID: 28320224; PubMed Central PMCID: PMCPMC5543416.

10. Insel T, Cuthbert B, Garvey M, Heinssen R, Pine DS, Quinn K, et al. Research domain criteria (RDoC): toward a new classification framework for research on mental disorders. Am J Psychiatry. 2010;167(7):748–51. Epub 2010/07/03. doi: 10.1176/appi.ajp.2010.09091379. PubMed PMID: 20595427.

11. Tamminga CA, Pearlson G, Keshavan M, Sweeney J, Clementz B, Thaker G. Bipolar and schizophrenia network for intermediate phenotypes: outcomes across the psychosis continuum. Schizophr Bull. 2014;40 Suppl 2(Suppl 2):S131-7. Epub 2014/02/25. doi: 10.1093/schbul/sbt179. PubMed PMID: 24562492; PubMed Central PMCID: PMCPMC3934403.

12. Karalunas SL, Fair D, Musser ED, Aykes K, Iyer SP, Nigg JT. Subtyping attention-deficit/hyperactivity disorder using temperament dimensions: toward biologically based nosologic criteria. JAMA Psychiatry. 2014;71(9):1015–24. Epub 2014/07/10. doi: 10.1001/jamapsychiatry.2014.763. PubMed PMID: 25006969; PubMed Central PMCID: PMCPMC4278404.

13. Li A, Zalesky A, Yue W, Howes O, Yan H, Liu Y, et al. A neuroimaging biomarker for striatal dysfunction in schizophrenia. Nat Med. 2020;26(4):558–65. Epub 2020/04/07. doi: 10.1038/s41591-020-0793-8. PubMed PMID: 32251404.

14. Yahata N, Morimoto J, Hashimoto R, Lisi G, Shibata K, Kawakubo Y, et al. A small number of abnormal brain connections predicts adult autism spectrum disorder. Nat Commun. 2016;7:11254. Epub 2016/04/15. doi: 10.1038/ncomms11254. PubMed PMID: 27075704; PubMed Central PMCID: PMCPMC4834637 a patent owned by Advanced Telecommunications Research (ATR) Institute International related to the present work [PCT/JP2014/061543 (WO2014178322)]. J.M., M.K., N.Y., R.H., K.S., T.W., Y.S., N.K. and K.K. are inventors of a patent owned by ATR Institute International related to the present work [PCT/JP2014/061544 (WO2014178323)]. G.L., J.M., M.K. and N.Y. are inventors of a patent application submitted by ATR Institute International related to the present work [JP2015-228970]. H.K. received honoraria for lectures related to the Japanese version of ADI-R from Kanekoshobo. M.Ku. was previously involved in the publication of the Japanese version of ADOS and ADI-R. M.Ku. also received honoraria for lectures related to the Japanese version of ADI-R and ADOS from Kanekoshobo. H.I. received honoraria for lectures by Shimadzu Co. and Nippon Boehringer Ingelheim Co., Ltd. H.T. received research grants from Takeda. H.T. has received honoraria for lectures by Otsuka, Meiji Seika, MSD, Dainippon-Sumitomo and GlaxoSmithKline. For the past three years, Y.O. declare the following potential conflicts of interest, although they are all unrelated to the current study. Y.O. has received honoraria for lectures by Otsuka, Dainippon Sumitomo, Astellas, Pfizer, Eli Lilly, Janssen, Meiji Seika Pharma, Mochida, Yoshitomi Yakuhin, Eizai and GlaxoSmithKline. For the past three years, K.K. declare the following potential conflicts of interest, although they are all unrelated to the current study. K.K. received research grants from Astellas, GlaxoSmithKline, Dainippon-Sumitomo, Eisai, MSD and Yoshitomi. K.K. has received honoraria for lectures by Daiichi-Sankyo, Otsuka, Meiji Seika, MSD, Astellas, Yoshitomi, Novartis, Eli Lilly, Dainippon-Sumitomo, Janssen, GlaxoSmithKline and Pfizer.

15. Zhang Y, Wu W, Toll RT, Naparstek S, Maron-Katz A, Watts M, et al. Identification of psychiatric disorder subtypes from functional connectivity patterns in resting-state electroencephalography. Nat Biomed Eng. 2021;5(4):309–23. Epub 2020/10/21. doi: 10.1038/s41551-020-00614-8. PubMed PMID: 33077939; PubMed Central PMCID: PMCPMC8053667.

16. Kaufmann T, Alnaes D, Doan NT, Brandt CL, Andreassen OA, Westlye LT. Delayed stabilization and individualization in connectome development are related to psychiatric disorders. Nat Neurosci. 2017;20(4):513–5. Epub 2017/02/22. doi: 10.1038/nn.4511. PubMed PMID: 28218917.

17. Finn ES, Shen X, Scheinost D, Rosenberg MD, Huang J, Chun MM, et al. Functional connectome fingerprinting: identifying individuals using patterns of brain connectivity. Nat Neurosci. 2015;18(11):1664–71. Epub 2015/10/13. doi: 10.1038/nn.4135. PubMed PMID: 26457551; PubMed Central PMCID: PMCPMC5008686.

18. Dinga R, Schmaal L, Penninx B, van Tol MJ, Veltman DJ, van Velzen L, et al. Evaluating the evidence for biotypes of depression: Methodological replication and extension of. Neuroimage Clin. 2019;22:101796. Epub 2019/04/03. doi: 10.1016/j.nicl.2019.101796. PubMed PMID: 30935858; PubMed Central PMCID: PMCPMC6543446.

19. He Y, Byrge L, Kennedy DP. Nonreplication of functional connectivity differences in autism spectrum disorder across multiple sites and denoising strategies. Hum Brain Mapp. 2020;41(5):1334–50. Epub 2020/01/10. doi: 10.1002/hbm.24879. PubMed PMID: 31916675; PubMed Central PMCID: PMCPMC7268009.

20. Yamashita A, Sakai Y, Yamada T, Yahata N, Kunimatsu A, Okada N, et al. Generalizable brain network markers of major depressive disorder across multiple imaging sites. PLoS Biol. 2020;18(12):e3000966. Epub 2020/12/08. doi: 10.1371/journal.pbio.3000966. PubMed PMID: 33284797; PubMed Central PMCID: PMCPMC7721148 following competing interests: M.K., N.Y., R.H., H.I., N.K. and K.K are inventors of a patent owned by Advanced Telecommunications Research (ATR) Institute International related to the present work [PCT/JP2014/061543 (WO2014178322)]. M.K., N.Y., R.H., N.K. and K.K. are inventors of a patent owned by ATR Institute International related to the present work [PCT/JP2014/061544 (WO2014178323)]. M.K. and N.Y. are inventors of a patent application submitted by ATR Institute International related to the present work [JP2015-228970]. A.Y. and M.K. are inventors of a patent application submitted by ATR Institute International related to the present work [JP2018-192842].

21. Feczko E, Miranda-Dominguez O, Marr M, Graham AM, Nigg JT, Fair DA. The Heterogeneity Problem: Approaches to Identify Psychiatric Subtypes. Trends Cogn Sci. 2019;23(7):584–601. Epub 2019/06/04. doi: 10.1016/j.tics.2019.03.009. PubMed PMID: 31153774; PubMed Central PMCID: PMCPMC6821457.

22. Xu K, Hu W, Leskovec J, Jegelka S. How powerful are graph neural networks? arXiv preprint arXiv:181000826. 2018.

23. Chang C-C, Lin C -J. LIBSVM: A library for support vector machines. ACM Trans Intell Syst Technol. 2011;2(3):Article 27. doi: 10.1145/1961189.1961199.

24. Van der Maaten L, Hinton G. Visualizing data using t-SNE. Journal of machine learning research. 2008;9(11).

25. Johnson WE, Li C, Rabinovic A. Adjusting batch effects in microarray expression data using empirical Bayes methods. Biostatistics. 2007;8(1):118–27. Epub 2006/04/25. doi: 10.1093/biostatistics/kxj037. PubMed PMID: 16632515.

26. Fortin JP, Cullen N, Sheline YI, Taylor WD, Aselcioglu I, Cook PA, et al. Harmonization of cortical thickness measurements across scanners and sites. Neuroimage. 2018;167:104–20. Epub 2017/11/21. doi: 10.1016/j.neuroimage.2017.11.024. PubMed PMID: 29155184; PubMed Central PMCID: PMCPMC5845848.

27. Fortin JP, Parker D, Tunç B, Watanabe T, Elliott MA, Ruparel K, et al. Harmonization of multi-site diffusion tensor imaging data. Neuroimage. 2017;161:149–70. Epub 2017/08/23. doi: 10.1016/j.neuroimage.2017.08.047. PubMed PMID: 28826946; PubMed Central PMCID: PMCPMC5736019.

28. Yu M, Linn KA, Cook PA, Phillips ML, McInnis M, Fava M, et al. Statistical harmonization corrects site effects in functional connectivity measurements from multi-site fMRI data. Hum Brain Mapp. 2018;39(11):4213–27. Epub 2018/07/03. doi: 10.1002/hbm.24241. PubMed PMID: 29962049; PubMed Central PMCID: PMCPMC6179920.

29. Rodriguez A, Laio A. Machine learning. Clustering by fast search and find of density peaks. Science. 2014;344(6191):1492–6. Epub 2014/06/28. doi: 10.1126/science.1242072. PubMed PMID: 24970081.

30. Sporns O, Chialvo DR, Kaiser M, Hilgetag CC. Organization, development and function of complex brain networks. Trends Cogn Sci. 2004;8(9):418–25. Epub 2004/09/08. doi: 10.1016/j.tics.2004.07.008. PubMed PMID: 15350243.

31. Bassett DS, Bullmore E. Small-world brain networks. Neuroscientist. 2006;12(6):512–23. Epub 2006/11/03. doi: 10.1177/1073858406293182. PubMed PMID: 17079517.

32. Reijneveld JC, Ponten SC, Berendse HW, Stam CJ. The application of graph theoretical analysis to complex networks in the brain. Clin Neurophysiol. 2007;118(11):2317–31. Epub 2007/09/29. doi: 10.1016/j.clinph.2007.08.010. PubMed PMID: 17900977.

33. Stam CJ, Reijneveld JC. Graph theoretical analysis of complex networks in the brain. Nonlinear Biomed Phys. 2007;1(1):3. Epub 2007/10/03. doi: 10.1186/1753-4631-1-3. PubMed PMID: 17908336; PubMed Central PMCID: PMCPMC1976403.

34. Guimerà R, Mossa S, Turtschi A, Amaral LA. The worldwide air transportation network: Anomalous centrality, community structure, and cities’ global roles. Proc Natl Acad Sci U S A. 2005;102(22):7794–9. Epub 2005/05/25. doi: 10.1073/pnas.0407994102. PubMed PMID: 15911778; PubMed Central PMCID: PMCPMC1142352.

35. Guimerà R, Nunes Amaral LA. Functional cartography of complex metabolic networks. Nature. 2005;433(7028):895–900. Epub 2005/02/25. doi: 10.1038/nature03288. PubMed PMID: 15729348; PubMed Central PMCID: PMCPMC2175124.

36. Girvan M, Newman ME. Community structure in social and biological networks. Proc Natl Acad Sci U S A. 2002;99(12):7821–6. Epub 2002/06/13. doi: 10.1073/pnas.122653799. PubMed PMID: 12060727; PubMed Central PMCID: PMCPMC122977.

37. Ying R, Bourgeois D, You J, Zitnik M, Leskovec J. Gnnexplainer: Generating explanations for graph neural networks. Advances in neural information processing systems. 2019;32:9240.

38. Plis SM, Hjelm DR, Salakhutdinov R, Allen EA, Bockholt HJ, Long JD, et al. Deep learning for neuroimaging: a validation study. Front Neurosci. 2014;8:229. Epub 2014/09/06. doi: 10.3389/fnins.2014.00229. PubMed PMID: 25191215; PubMed Central PMCID: PMCPMC4138493.

39. Yan W, Calhoun V, Song M, Cui Y, Yan H, Liu S, et al. Discriminating schizophrenia using recurrent neural network applied on time courses of multi-site FMRI data. EBioMedicine. 2019;47:543–52. Epub 2019/08/20. doi: 10.1016/j.ebiom.2019.08.023. PubMed PMID: 31420302; PubMed Central PMCID: PMCPMC6796503.

40. Sui J, Jiang R, Bustillo J, Calhoun V. Neuroimaging-based Individualized Prediction of Cognition and Behavior for Mental Disorders and Health: Methods and Promises. Biol Psychiatry. 2020;88(11):818–28. Epub 2020/04/28. doi: 10.1016/j.biopsych.2020.02.016. PubMed PMID: 32336400; PubMed Central PMCID: PMCPMC7483317.

41. Rakhimberdina Z, Liu X, Murata AT. Population Graph-Based Multi-Model Ensemble Method for Diagnosing Autism Spectrum Disorder. Sensors (Basel). 2020;20(21). Epub 2020/10/28. doi: 10.3390/s20216001. PubMed PMID: 33105909; PubMed Central PMCID: PMCPMC7660214.

42. Rakhimberdina Z, Murata T, editors. Linear Graph Convolutional Model for Diagnosing Brain Disorders2020; Cham: Springer International Publishing.

43. Yao D, Liu M, Wang M, Lian C, Wei J, Sun L, et al., editors. Triplet Graph Convolutional Network for Multi-scale Analysis of Functional Connectivity Using Functional MRI2019; Cham: Springer International Publishing.

44. Yang H, Li X, Wu Y, Li S, Lu S, Duncan JS, et al., editors. Interpretable Multimodality Embedding of Cerebral Cortex Using Attention Graph Network for Identifying Bipolar Disorder2019; Cham: Springer International Publishing.

45. Qiu S, Joshi PS, Miller MI, Xue C, Zhou X, Karjadi C, et al. Development and validation of an interpretable deep learning framework for Alzheimer’s disease classification. Brain. 2020;143(6):1920-33. Epub 2020/05/02. doi: 10.1093/brain/awaa137. PubMed PMID: 32357201; PubMed Central PMCID: PMCPMC7296847.

46. Castelvecchi D. Can we open the black box of AI? Nature. 2016;538(7623):20-3. Epub 2016/10/07. doi: 10.1038/538020a. PubMed PMID: 27708329.

47. Huang ZA, Zhu Z, Yau CH, Tan KC. Identifying Autism Spectrum Disorder From Resting-State fMRI Using Deep Belief Network. IEEE Trans Neural Netw Learn Syst. 2021;32(7):2847–61. Epub 2020/07/22. doi: 10.1109/tnnls.2020.3007943. PubMed PMID: 32692687.

48. Heinsfeld AS, Franco AR, Craddock RC, Buchweitz A, Meneguzzi F. Identification of autism spectrum disorder using deep learning and the ABIDE dataset. Neuroimage Clin. 2018;17:16–23. Epub 2017/10/17. doi: 10.1016/j.nicl.2017.08.017. PubMed PMID: 29034163; PubMed Central PMCID: PMCPMC5635344.

49. Wang Y, Tang S, Zhang L, Bu X, Lu L, Li H, et al. Data-driven clustering differentiates subtypes of major depressive disorder with distinct brain connectivity and symptom features. The British Journal of Psychiatry. 2021;219(5):606–13. Epub 2021/07/30. doi: 10.1192/bjp.2021.103.

50. Yan CG, Chen X, Li L, Castellanos FX, Bai TJ, Bo QJ, et al. Reduced default mode network functional connectivity in patients with recurrent major depressive disorder. Proc Natl Acad Sci U S A. 2019;116(18):9078–83. Epub 2019/04/14. doi: 10.1073/pnas.1900390116. PubMed PMID: 30979801; PubMed Central PMCID: PMCPMC6500168.

51. Tzourio-Mazoyer N, Landeau B, Papathanassiou D, Crivello F, Etard O, Delcroix N, et al. Automated anatomical labeling of activations in SPM using a macroscopic anatomical parcellation of the MNI MRI single-subject brain. Neuroimage. 2002;15(1):273–89. Epub 2002/01/05. doi: 10.1006/nimg.2001.0978. PubMed PMID: 11771995.

52. Rosenthal G, Váša F, Griffa A, Hagmann P, Amico E, Goñi J, et al. Mapping higher-order relations between brain structure and function with embedded vector representations of connectomes. Nat Commun. 2018;9(1):2178. Epub 2018/06/07. doi: 10.1038/s41467-018-04614-w. PubMed PMID: 29872218; PubMed Central PMCID: PMCPMC5988787.

53. Park H-S, Jun C-H. A simple and fast algorithm for K-medoids clustering. Expert Systems with Applications. 2009;36(2, Part 2):3336-41. doi: https://doi.org/10.1016/j.eswa.2008.01.039.

